# Mental fatigue has only marginal effects on static balance control in healthy young adults

**DOI:** 10.1101/2023.07.05.547754

**Authors:** Kerstin Weissinger, Margit Midtgaard Bach, Anna Brachmann, John F. Stins, Peter Jan Beek

## Abstract

We examined the influence of mental fatigue on static balance control in healthy young adults to gain greater clarity about this issue than provided in previous research. Based on the prevailing assumption in pertinent literature, we hypothesized that mental fatigue leads to a reduced cognitive regulation of quiet upright standing, as reflected in center of pressure (COP) excursions. More specifically, we hypothesized that the influence of mental fatigue on balance control depends on the attentional effort required by the balance tasks being performed. To test these hypotheses, 44 young adults (24 women and 20 men) were quasi-randomly assigned to either an experimental group that was mentally fatigued (using the TloadDback-task with individualized settings) or a control group (who watched a documentary). Before and after the intervention the participants performed six balance tasks that differed in (attentional) control requirements, while their COP was being recorded. From these time-series sway variability, mean speed, and sample entropy were calculated and analyzed statistically. Additionally, mental fatigue was assessed using VAS scales. Statistical analyses confirmed that the balance tasks differed in control characteristics and that mental fatigue was elevated in the experimental group, but not in the control group. Nevertheless, no significant main effects of mental fatigue were found on any of the COP measures of interest, except for some non-robust and difficult to interpret interaction effects involving the factor group. These results suggest that, in young adults, postural control in static balance tasks is largely automatic and unaffected by mental fatigue.

## 1. Introduction

Mental fatigue (MF) is a common psychophysiological phenomenon that occurs during prolonged demanding cognitive activities (Boksem, Meijman, & Lorist, 2005; Hockey, 2013; Van Cutsem, et al., 2017). It results from overusing the brain’s capacity and resources (Tanaka, 2015), and has various manifestations, including feeling tired, worn out or lethargic, and having headaches (Hockey, 2013). Due to its impact on cognitive performance, MF typically has adverse effects on the performance of daily activities that require cognitive resources, making it less effective and efficient. In pertinent experimental studies, MF has been found to interfere with both executive and attentional functions (Borragan, Slama, Bartolomei, & Peigneux, 2017; Tanaka, 2015), as evidenced by longer times needed to process, plan, and respond to stimuli. MF also degrades the accuracy of responses in performing cognitive tasks, such as the Stroop task, the psychomotor vigilance task and the AX-continuous performance task (Hachard, Noé, Ceyte, Trajin, & Paillard, 2020). Furthermore, MF makes it harder to identify and focus on relevant information and suppress irrelevant stimuli (Borragan, et al., 2017), suggesting that MF induces attention shifts from being goal-directed towards stimulus driven (Boksem, et al., 2005). Finally, MF has been shown to interfere with motor performance. For example, Lew and Qu (2014) found that mentally fatigued young adults displayed an increased risk of slipping when exposed to laboratory-induced slip-like perturbations. Based on such findings, it has been suggested that MF poses a risk for postural stability, potentially leading to accidents (Lew & Qu, 2014; Tanaka, 2015).

In this study we focus on one instance of postural stability, namely quiet upright standing. This is a fundamental motor skill that is essential for performing a broad variety of daily life activities in a safe and efficient manner (Hachard, et al., 2020). For decades it has been assumed that maintaining stability during unperturbed upright standing is an automatic and largely reflex-driven process, which thus requires little or no attentional regulation (Kerr, Condon, & McDonald, 1985; Nashner, 1976; Takakusaki, Saitoh, Harada, & Kashiwayanagi, 2004; Teasdale, Bard, LaRue, & Fleury, 1993). However, under certain experimental conditions, such as cognitive-postural dual-tasks and sensory manipulations (e.g., maintaining balance with eyes closed), it can be observed that attentional resources are tapped, even in simple postural tasks like stable stance (Papegaaij, Taube, Baudry, Otten, & Hortobagyi, 2014; Ruffieux, Keller, Lauber, & Taube, 2015; Teo, Goodwill, Hendy, Muthalib, & Macpherson, 2018). In such instances, an increase in cortical activation has been observed compared to performing the same task without visual or proprioceptive manipulations or having to perform a concurrent cognitive task (Bergamin, et al., 2014; Prado, Stoffregen, & Duarte, 2007; Teo, et al., 2018). This increase in cortical activity might reflect elevated attentional demands to ensure postural stability (Lajoie, Teasdale, Bard, & Fleury, 1993; Papegaaij, et al., 2014). Another observation pointing towards a possible link between postural stability and cognitive resources is that older individuals or individuals with mild cognitive impairment are less stable during quiet stance and exhibit a slower gait speed compared to their cognitively fitter counterparts (Behrens, et al., 2017; Deschamps, Beauchet, Annweiler, Cornu, & Mignardot, 2014; Grobe, et al., 2017; Muir, et al., 2012). Both observations raise the question whether MF, which is assumed to deplete cortical resources (Tanaka, 2015), adversely influences the performance of static balance tasks.

Last year, two systematic reviews were published that examined the influence of MF on balance in healthy individuals (Brahms, et al., 2022; Pitts & Bhatt, 2023), with partially overlapping citations but no reference to each other. Three important observations can be gleaned from those reviews. First, the literature is rather limited in size and consists of circa 10 experimental studies (depending on the adopted inclusion and exclusion criteria). The small body of literature makes the research field easy to oversee, but also highlights the need to broaden the database to be able to draw solid, reliable conclusions. Second, both reviews, as well as the papers cited therein, work from the assumption that MF specifically influences the cortical regulation of balance, with little or no effect on subcortical and spinal control loops. Therefore, effects of MF are expected to occur when the balance task requires greater cognitive involvement, as in challenging postural situations, and/or when the automatic regulation of balance is hampered due to pathology. The question is whether these expectations hold true. However, as it stands, and this is the third noteworthy observation, the evidence for a robust influence of MF on posture is inconclusive, as is apparent from the upshot of both systematic reviews. Whereas Brahms, et al. (2022) concluded unequivocally that MF can adversely affect balance in healthy young adults, Pitts and Bhatt (2023) came to a more nuanced conclusion as some studies only showed modest, partial or even null effects (Varas-Diaz, Kannan, & Bhatt, 2020). Differences in research findings might have been due to various methodological factors, including the method(s) used to induce MF, the type of balance control investigated (e.g., volitional versus reactive), the experimental task used, the balance measures employed, the study population (e.g., healthy young adults, healthy old adults, individuals with stroke) and the number of participants (and thus statistical power).

The present study was conducted to help resolve the existing ambiguity in study results by assessing the effect of MF on static balance control in a rigorously controlled, well-powered experiment, using multiple balance measures, thus extending the data base of experimental findings for drawing solid conclusions and making theoretical inferences. A crucial factor in achieving this aim resides in the selection and appropriation of a suitable method to induce MF in a laboratory setting. Various methods have been employed for this purpose, which differ considerably in terms of task requirements and duration. However, most MF studies published to date used a prolonged computerized task to induce MF, based on the notion that MF needs extended time to accumulate. Examples are the 90-minute AX-CPT (Hachard, et al., 2020; Noé, Hachard, Ceyte, Bru, & Paillard, 2021), the Stroop Task performed for either 90 (Tassignon, et al., 2020; Verschueren, et al., 2020) or 45 minutes (Schouppe, et al., 2019), the 30-minute Psychomotor Vigilance Task (Deschamps, Magnard, & Cornu, 2013), and the 60-minute Stop-Signal Task (Behrens, et al., 2017; Varas-Diaz, et al., 2020). Although all these tasks reliably induced MF on a group level, they did not take individual differences in susceptibility to MF into account. The same task may thus have resulted in different MF levels across participants, making it difficult to draw valid and reliable conclusions about the influence of MF on motor performance, including postural control (Noé, et al., 2021). Since previous research has revealed large individual differences in susceptibility to MF (O’Keeffe, Hodder, & Lloyd, 2019), we chose the TloadDback task for inducing MF (Borragan, et al., 2017; Borragan, Slama, Destrebecqz, & Peigneux, 2016) to examine its effect on posture, because this task has the advantage that its settings can be adapted to an individually predefined maximum cognitive load, thus rendering the degree of MF equivalent across participants. Moreover, it has been demonstrated that, with this task, only 16 minutes are required to induce a sufficient level of MF (O’Keeffe, et al., 2019).

In the present study we examined the effects of MF on postural stability in healthy young adults. We chose this population because we wished to determine how generic a possible effect of MF on static posture would be. Patently, if such an effect would exist in young adults, it would most likely also exist in other physically and mentally less prone populations. In accordance with previous findings and the assumption of impaired cognitive control, our main hypothesis was that MF has an adverse effect on balance control during quiet upright standing, as evidenced by changes in the characteristics of the center-of-pressure (COP) recordings. Specifically, we expected the sway variability (as indexed by the standard deviation) and the speed of the COP to increase, and the degree of randomness (as indexed by the sample entropy) in the COP time-series to decrease with MF. All these measures, including the sample entropy, have been used in previous studies (Becker & Hung, 2020; Donker, Roerdink, Greven, & Beek, 2007; Potvin-Desrochers, Richer, & Lajoie, 2017; Richer & Lajoie, 2020; Roerdink, et al., 2006; J. F. Stins, Michielsen, Roerdink, & Beek, 2009) on postural control during standing tasks, thus allowing comparison of findings across studies. Furthermore, we hypothesized that the influence of MF on balance control depends on the attentional effort required by the balance tasks being performed. To this end, we included several static balance tasks in the experimental design that differed in complexity and attentional demands. We combined two postures (hip-broad and tandem stance) with three task manipulations (eyes open, eyes closed, and dual task), for a total of six balance tasks. Balance control was hypothesized to be less stable and more challenging, requiring greater attentional control, in tandem stance compared to hip-broad stance, as well as with eyes closed compared to eyes open, while the addition of a cognitive dual task was expected to reduce attentional control over the balance task being performed.

To test our research hypotheses, we first verified that the postural manipulations indeed resulted in different COP patterns in the absence of MF. After all, this must be the case to examine the hypothesis that the influence of MF increases with task complexity. In particular, based on previous studies, we expected that tandem stance is less stable than hip-broad stance, that standing with eyes closed is less stable than with eyes open, as reflected in increased postural sway and reduced COP complexity in both comparisons, and that adding a cognitive dual task leads to reduced postural sway and increased COP complexity, presumably due to a reduction in attentional focus (Becker & Hung, 2020; Donker, et al., 2007; Potvin-Desrochers, et al., 2017; Rhea, Diekfuss, Fairbrother, & Raisbeck, 2019; Yamada & Raisbeck, 2021). These expectations were verified by examining the COP fluctuations prior to the MF intervention. The protocol of this study, together with the hypotheses and statistical analyses, were pre-registered on Open Source Framework (OSF): https://osf.io/2e9gs.

## 2. Methods

### 2.1. Participants

The sample size required for this study was calculated using G*Power (Faul, Erdfelder, Lang, & Buchner, 2007). For a statistical power of 0.95, a standard alpha error probability of 0.01 and a medium effect size of f = 0.25, 36 participants would be required. To remain on the conservative side and anticipate the possibility of participant and/or data loss, a convenience sample of 44 young adults were recruited for the study in the immediate vicinity of the investigators. As a result, the sample mostly comprised students and employees of the Vrije Universiteit Amsterdam. Participants had to be between 20 and 35 years old and healthy, without regard of sex. All participants (24 female and 20 male, average age 25.86 ± 3.26 years; mean ± standard deviation [SD]) signed an informed consent form prior to the experiment. They were asked not to engage in vigorous exercise 48 hours before the experiment(Deschamps, et al., 2013) and to consume no beverages containing alcohol, caffeine, or other stimulants (Hachard, et al., 2020) for minimally two hours before the experiment. Participants were further requested to avoid sleep deprivation as much as possible the night before the experiment. As sleep quantity and quality can interfere with both cognitive and motor performance, participants were asked about their sleep before the experiment by posing two questions from the ‘Pittsburg Sleep Quality Index’ (Buysse, Reynolds, Monk, Berman, & Kupfer, 1989), namely (1) How many hours of actual sleep did you get at night? (This may be different than the number of hours spent in bed.), and (2) How would you rate your sleep quality? (1 = very good, 2 = fairly good, 3 = fairly bad, 4 = bad). The same requirements had to be fulfilled before the familiarization session, in which the stimulus duration time was determined (see below).

### 2.2. Protocol

The experimental protocol is shown schematically in Fig. 1. The protocol fully complied with the Declaration of Helsinki and was approved by the local ethics committee of the Faculty of Behavioral and Movement Sciences of the Vrije Universiteit Amsterdam (VCWE-2022-154R2).

**Fig. 1.**
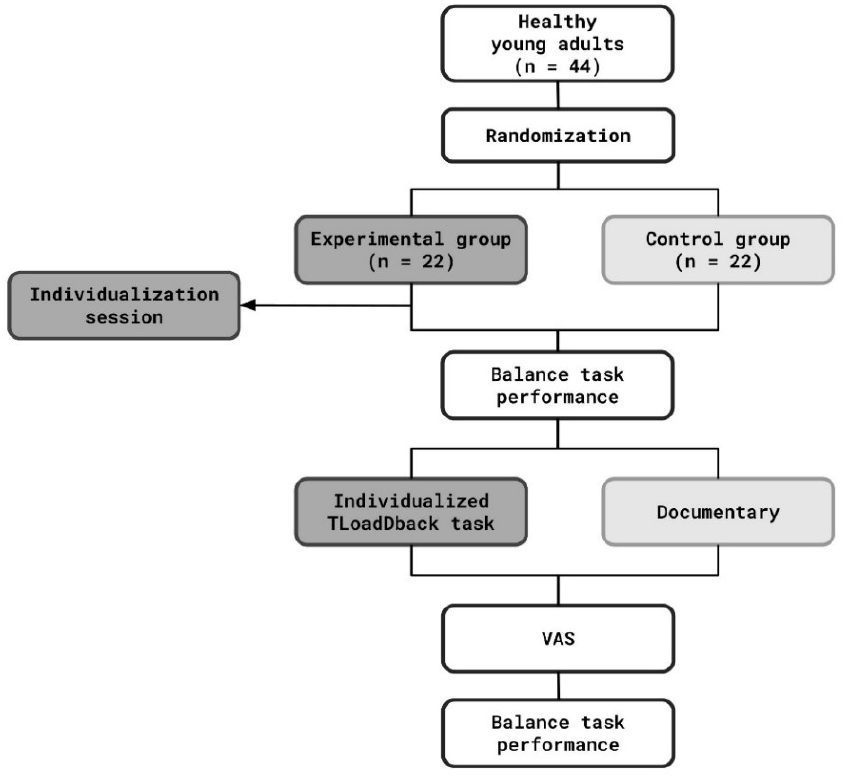
Schematic representation of the experimental protocol.

To assess the effect of MF on postural stability, a mixed design was used. Participants were quasi-randomly (i.e., alternately) assigned to either the experimental or control group. The participants in the experimental group were mentally fatigued using the TloadDback task, whereas the participants in the control group watched a neutral documentary of equal duration. The TloadDback task was adapted to the cognitive capacity of each individual participant in the experimental group. To this end, the participants in this group underwent an extra measurement session.

The assessment of postural balance before and after the intervention consisted of six balance tasks (two postures under three conditions); all of which were performed on a force plate. To avoid an order effect and consequent potential confounding of the results, the six balance tasks were performed in randomized order. Randomization was achieved with the help of a web application available at http://random.org. Right before the pre-intervention balance tests, all participants were allowed a few minutes to familiarize themselves with the experimental tasks and setup until they felt comfortable with them. Right before the post-intervention balance tests, subjective levels of MF were evaluated using visual analog scales (VAS).

If a participant was unable to maintain a certain balance task for the full recording time, the trial in question was repeated immediately. Participants were allowed a maximum of three attempts to successfully complete any given balance task. If all three attempts were unsuccessful, or if recordings turned out to be invalid, the trials were excluded for that balance task of the respective participant, while successful recordings of the other tasks were still included in the analysis.

### 2.3. Interventions

The Time load Dual-back (TloadDback) developed by Borragán et al. (2016, 2017) combines two information processing tasks, the N-back working-memory updating task and a parity task, the odd/even decision task. During the TloadDback task, letters and digits appear alternately (letter/digit/letter/…) on a computer screen. The participant was instructed to press “2” for even numbers and “3” for odd numbers on a numeric keyboard using their right hand. Furthermore, they should indicate whether the depicted letter equals the letter that was presented N letters back (with N = 1) by pressing the space bar with their left hand.

To induce similar levels of MF in all participants the task was adapted such that it imposed the same cognitive load on all participants. Instead of adapting N as in an N-back task, this was done by adapting the duration for which each stimulus was displayed on the screen, termed the Stimulus Time Duration (STD). If the task is being sped up by gradually reducing the STD, more and more effort is required to maintain accurate performance, thus arguably depleting cognitive resources, and increasing MF (Barrouillet, Bernardin, Portrat, Vergauwe, & Camos, 2007). Following the recommendations of (Borragan, et al., 2017), the STD was reduced until an accuracy of > 85 % could no longer be sustained. The minimal possible STD of each participant in the experimental group was determined in a familiarization session held 1-7 days prior to the actual experiment. To minimize the effect of day-to-day differences, the actual experimental session was scheduled at the same time of day as the first session. This was accomplished (±1 h) for all except two participants (+ 4 h / -2.5 h compared to the first session).

For the control intervention a 16-minute excerpt of the beginning of the BBC documentary “Earth” (Fothergill and Linfield 2007) was chosen as this control task was also used in previous studies, thus allowing comparison of results (Hachard, et al., 2020; Varas-Diaz, et al., 2020).

### 2.4. Assessment of subjective MF

Subjective levels of MF were assessed with a digital visual analog scale (VAS) using the AVAS-Software Marsh-Richard et al. 2009. To avoid response shift bias (Howard 1980), a retrospective pretest design was used in which the participants only performed the VAS after the intervention (Gorrall, Curtis, Little, & Panko, 2016). Participants were asked to rate their MF on the VAS in both an absolute and a relative manner. Directly after the intervention (TloadDback task or documentary, respectively), they were asked to rate MF on a scale ranging from ‘not at all fatigued’ (≜0) to ‘extremely fatigued’ (≜100) (absolute score). Subsequently, they were asked to reflect and compare their degree of MF after to before the intervention, on a scale ranging from ‘much less fatigued’ (≜-50) to ‘much more fatigued’ (≜50), with 0 indicating no change (relative score).

### 2.5. Assessment of postural control

Postural control was assessed in two static balance postures: (1) hip-broad stance, with feet positioned hip broad and parallel to each other, toes pointing forward, and (2) tandem stance, with one foot placed directly in front of the other with the heel of the front foot touching the big toe of the back foot. No instructions were given about which foot to place in front. Both postures were carried out in three conditions: (i) eyes open, (ii) eyes closed, and (iii) dual task (balance task combined with a cognitive task). The cognitive task consisted of counting backwards in steps of 7 from a randomly given number between 150 and 300. The subtractions were carried out silently, i.e., without vocalization (John F. Stins, Roerdink, & Beek, 2011). Participants were instructed to prioritize the balance task over the cognitive task. The performance of the cognitive task was not further analyzed.

Postural sway was recorded with a (1 x 1 m) custom-made strain gauge force plate, consisting of eight force sensors, four of which measured the forces on the *z*-axis (vertical), two the forces on the *x*-axis, and two the forces on the *y*-axis. The resulting eight signals were automatically converted by the program underlying the force plate to a COP time-series in the medio-lateral (ML) and anterior-posterior (AP) direction. The sampling rate of the force plate was 100 Hz. The recording time was set to 60 seconds, since recording times of 60 seconds and longer are known to be beneficial to detect differences between groups when calculating sample entropy (Montesinos et al. 2018). Participants were instructed to maintain each posture for 60 seconds and sway as little as possible. In the balance tasks with eyes open (including the dual-task conditions), participants were asked to focus on a circle attached at eye-height two meters in front of them (J. F. Stins, et al., 2009).

The recorded data were analyzed in MATLAB (Mathworks, Inc., Version R2022a). First, the COP time-series were processed and low-pass filtered at 12.5 Hz using a bi-directionally second-order Butterworth to reduce signal noise. Subsequently, the COP time-series were analyzed in the anteroposterior (AP) and mediolateral (ML) directions separately by calculating three outcome measures: sway variability (standard deviation), mean speed, and degree of randomness (sample entropy).

#### Sway variability

The standard deviation was calculated as a measure of sway variability, in accordance with its definition:

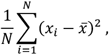

where *N*is the number of samples, *x*_*i*_ the displacement at time *t*_*i*_, and 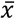 the time-series mean.

#### Mean speed

The mean displacement speed was calculated as the mean of the absolute values of the difference quotients of the displacements for the time-series in question:

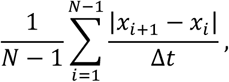

where *N*is the number of samples, *x*_*i*_ the displacement at time *t*_*i*_, and Δ*t* the time window between two samples. Since the mean speed is independent of the sampling time, it allows for a clean comparison across studies.

#### Sample entropy

Richman and Moorman (2000) defined the sample entropy in terms of conditional probabilities of similar sequences. Sample entropy *SampEn*(*m, r, N*) is the negative value of the natural logarithm of similarity counts of *m* + 1 points (*A*) over similarity counts of *m* points (*B*) among all points (*N*) of the sequence, where similarities are reached when distances between sample vectors are below a cut-off radius *r*:

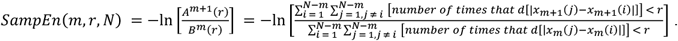

It is well known that the calculation of sample entropy is rather sensitive to the number of points/template length *m*, similarity tolerance *r*, and the total data number of data points *N*. We therefore calculated the sample entropy using two different parameter settings, based on two methods, which have both been recommended in the literature:

1. Lake, Richman, Griffin, and Moorman (2002) provide a method to select parameters based on minimizing the maximum relative error of sample entropy, which resulted in *m* = 3, *r* = 0.05, and *N* = 6000. This method is well established and commonly used in the analysis of physiological signals.
2. Montesinos, Castaldo, and Pecchia (2018) developed recommendation for parameter settings specifically for the analysis of COP time-series, which are *m* = 4, *r* = 0.25, and *N* = 6000.

Sample entropy values were calculated in MATLAB using a script by Martínez-Cagigal (2018).

## 3. Statistical analysis

First, all COP data (i.e., pre- and post-intervention) were analyzed for completeness and abnormalities, such as outliers. To reduce the impact of these outliers on the statistical analysis the ‘winsorizing’ method was applied to each of the three outcome measures. This data-clipping method replaces the score of an outlier with a determined minimum/maximum (Field, 2018). Specifically, if, *x* < *mean* − 2 · *SD* or x > *mean* + 2 · *SD, x* was replaced by the minimum or maximum value (*mean* − 2 · *SD, mean* + 2 · *SD*, respectively). The winsorized COP outcome measures were used in the subsequent analyses.

Two sets of repeated-measures ANOVAs with a mixed 2 (group) x 2 (posture) x 3 (condition) design were performed, followed by post-hoc tests for significant results in the form of pair-wise comparisons with Bonferroni correction.

The first set of ANOVAs examined whether the postural control during the six balance tasks indeed differed in the expected manner by comparing the COP outcome measures across those tasks. The factor group was added in this analysis to evaluate possible differences in this regard between the experimental and control group before the intervention (i.e., at baseline). The second set of ANOVAs assessed the influence of MF on postural stability by comparing the effect of the interventions on the COP outcome measures for different postures and conditions, i.e., across the six balance tasks. For this purpose, relative symmetric change scores for all three COP measures were calculated as follows:

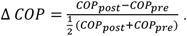

Dividing the absolute change score by the arithmetic mean is a standard procedure for data normalization to account for fluctuations in a pair of variables, provided that both are larger than zero (Burr & Nesselroade, 1990; Vartia, 1976).

The resulting *ΔCOP* measures were analyzed by conducting a repeated-measures ANOVA with the described design for each COP measure separately. In addition to the main effects, interaction effects were examined by means of post-hoc-tests with a Bonferroni correction. Effect sizes are reported as partial eta squared (η_p_^2^) for all main and interaction effects. By convention, 0.01 represents a small effect, 0.06 a medium effect, and 0.14 a large effect (Cohen, 1988).

Before conducting the ANOVAs, we assessed whether the data met the assumption of normality and sphericity (*ε*) by carrying out Kolmogorov-Smirnov (K-S) tests with Lilliefors correction and Mauchly’s test of sphericity. If the sphericity assumption was violated, the Greenhouse-Geisser correction was applied (Field 2018). Sleep quality and quantity, as well as the subjectively rated levels of induced mental fatigue, were assessed by means of Mann-Whitney tests. For all tests the level of significance was set at *p* < 0.05. All statistical analyses were performed in SPSS Statistics for Windows, version 27 (SPSS Inc., Chicago, IL., USA). Means and standard errors are reported in the statistical results unless specified otherwise.

## 4. Results

### 4.1. Participants

All participants except one male participant from the experimental group were included in the analysis. The participant in question was excluded because he had poor balance control and was unable to safely perform any balance tasks. After the participant’s exclusion, the experimental and control group remained similar in terms of sex and age distribution. The experimental group consisted of 11 female and 10 male participants with a mean age of 25.6 ± 3.8 years, while the control group consisted of 13 female and 9 male participants with a mean age of 26.1 ± 2.7 years.

### 4.2. Initial analysis of the COP outcome measures

An initial analysis of all pre and post COP outcome measures revealed that a few of them contained extreme values (46 out of 1032 investigated variables), which were adjusted by means of winsorizing. The Kolmogorov-Smirnov (K-S) tests with Lilliefors correction showed that the COP data were distributed normally. The Mauchly’s test of sphericity indicated a departure from sphericity for some variables, following which the Greenhouse-Geisser correction was applied.

### 4.3. COP behavior before intervention

As expected, the manipulation of the balance tasks had marked effects on the COP behavior of the participants. This can be seen in Fig. 2, which shows representative COP trajectories from a single participant for all six balance tasks before intervention.

**Fig. 2.**
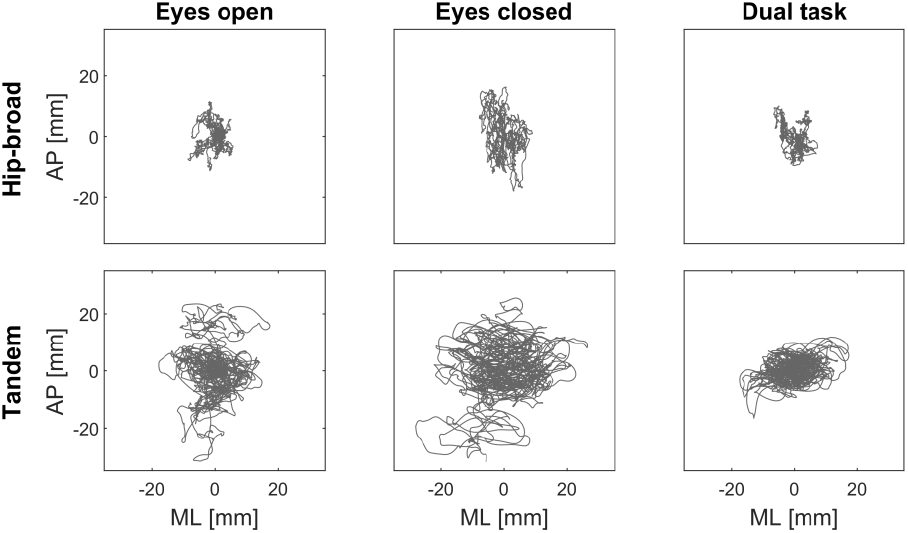
COP trajectories of a representative participant for the six balance tasks before intervention.

The effectiveness of the balance task manipulation was also evident from the results of the 2 x 2 x 3 repeated-measures ANOVAs that were performed on the pre-intervention COP outcome measures. These results are collected in Table 1 and Table 2.

**Table 1.** Statistical results of the repeated-measures ANOVA on the pre-intervention sway variability and mean speed values; *df* = degree of freedom, * = significant

| Effects |  | Sway variability |  |  |  | Mean speed |  |  |  |
| --- | --- | --- | --- | --- | --- | --- | --- | --- | --- |
| | | *df* (1,2) | *F* | *p* | $\eta_p^2$ | *df* (1,2) | *F* | *p* | $\eta_p^2$ |
| Group | ML | 1,41 | 3.24 | 0.079 | 0.07 | 1,41 | 2.80 | 0.102 | 0.06 |
|  | AP | 1,41 | 0.12 | 0.734 | 0.01 | 1,41 | 0.97 | 0.329 | 0.02 |
| Posture | ML | 1,41 | 722.13 | **<0.001\*** | 0.95 | 1,41 | 445.25 | **<0.001\*** | 0.92 |
|  | AP | 1,41 | 101.00 | **<0.001\*** | 0.71 | 1,41 | 285.88 | **<0.001\*** | 0.88 |
| Condition | ML | 1.2,51.15 | 165.81 | **<0.001\*** | 0.80 | 1.2,48.1 | 239.60 | **<0.001\*** | 0.85 |
|  | AP | 1.8,72.16 | 34.33 | **<0.001\*** | 0.46 | 1.2,46.9 | 167.54 | **<0.001\*** | 0.80 |
| Posture x Group | ML | 1,41 | 0.46 | 0.502 | 0.01 | 1,41 | 2.96 | 0.093 | 0.07 |
|  | AP | 1,41 | 0.76 | 0.382 | 0.02 | 1,41 | 0.55 | 0.463 | 0.01 |
| Condition x Group | ML | 1.2,51.15 | 3.03 | 0.079 | 0.07 | 1.2,48.1 | 0.57 | 0.481 | 0.01 |
|  | AP | 1.8,72.16 | 0.28 | 0.731 | 0.01 | 1.2,46.9 | 0.75 | 0.409 | 0.02 |
| Posture x Condition | ML | 1.3,53.7 | 185.89 | **<0.001\*** | 0.82 | 1.2,48.4 | 218.74 | **<0.001\*** | 0.84 |
|  | AP | 1.7,71.6 | 16.26 | **<0.001\*** | 0.28 | 1.1,46.1 | 106.91 | **<0.001\*** | 0.72 |
| Posture x Condition x Group | ML | 1.3,53.7 | 1.03 | 0.336 | 0.02 | 1.2,48.4 | 0.75 | 0.413 | 0.02 |
|  | AP | 1.7,71.6 | 0.59 | 0.537 | 0.01 | 1.1,46.1 | 0.85 | 0.702 | 0.01 |

**Table 2.** Statistical results of the repeated-measures ANOVA on the pre-intervention sample entropy values; right: calculated using Lake et al.’s method; left: calculated using Montesinos et al.et al.’s method; *df* = degree of freedom, * = significant

| Effects |  | Sample entropy: m = 3, r = 0.05 |  |  |  | Sample entropy: m = 4, r = 0.25 |  |  |  |
| --- | --- | --- | --- | --- | --- | --- | --- | --- | --- |
| | | *df* (1,2) | *F* | *p* | $\eta_p^2$ | *df* (1,2) | *F* | *p* | $\eta_p^2$ |
| Group | ML | 1,41 | 2.15 | 0.151 | 0.05 | 1,41 | 2.19 | 0.147 | 0.05 |
|  | AP | 1,41 | 0.23 | 0.632 | 0.01 | 1,41 | 0.07 | 0.787 | 0.01 |
| Posture | ML | 1,41 | 236.25 | **<0.001\*** | 0.85 | 1,41 | 15.55 | **<0.001\*** | 0.28 |
|  | AP | 1,41 | 0.02 | 0.967 | 0.00 | 1,41 | 31.51 | **<0.001\*** | 0.44 |
| Condition | ML | 1.8,73.4 | 7.01 | **0.002\*** | 0.16 | 2,82 | 3.20 | **0.046\*** | 0.07 |
|  | AP | 1.8,73.8 | 8.89 | **0.001\*** | 0.18 | 2,82 | 5.22 | **0.007\*** | 0.11 |
| Posture x Group | ML | 1,41 | 3.06 | 0.088 | 0.97 | 1,41 | 2.40 | 0.129 | 0.06 |
|  | AP | 1,41 | 1.50 | 0.227 | 0.04 | 1,41 | 1.03 | 0.317 | 0.02 |
| Condition x Group | ML | 1.8,73.4 | 2.08 | 0.137 | 0.05 | 2,82 | 1.94 | 0.150 | 0.05 |
|  | AP | 1.8,73.8 | 1.37 | 0.259 | 0.03 | 2,82 | 0.93 | 0.397 | 0.02 |
| Posture x Condition | ML | 1.8,73.9 | 2.64 | 0.084 | 0.06 | 2,82 | 2.71 | 0.073 | 0.06 |
|  | AP | 1.9,78.8 | 1.30 | 0.277 | 0.03 | 2,82 | 1.50 | 0.229 | 0.04 |
| Posture x Condition<br>x Group | ML | 1.8,73.9 | 1.92 | 0.158 | 0.05 | 2,82 | 1.89 | 0.160 | 0.04 |
|  | AP | 1.9,78.8 | 2.93 | 0.062 | 0.00 | 2,82 | 1.04 | 0.357 | 0.03 |

Three noteworthy sets of findings can be gleaned from these tables in view of our expectations regarding the balance task manipulation. First, as expected, significant main effects of posture occurred. The tandem stance proved more difficult to maintain than the hip-broad stance, as evidenced by significantly larger sway variability and higher speeds in both the ML- and AP-direction.

Posture also had a significant effect on the sample entropy in both directions when using Montesinos et al.’s method, but only in the ML-direction when using Lake et al.’s method. The sample entropy values were significantly lower for the tandem than the hip-broad stance, indicating tighter postural control.

Second, significant main effects of condition were found for all three COP outcome measures. Post-hoc analyses with Bonferroni correction for these effects revealed that sway variability and mean speed values were significantly higher when standing with closed eyes compared to standing with eyes open or adding a cognitive dual-task. These results confirmed our expectation that postural control is more difficult and hence less stable with eyes closed. Furthermore, for both calculation methods the sample entropy measures were significantly higher in the dual-task condition compared to standing with either eyes open or closed (without dual-task).

Third, significant ‘posture x condition’ interactions were found for sway variability and mean speed in both directions, but not for sample entropy, regardless of the method used for its calculation. Post-hoc tests revealed that the effect of posture was amplified with eyes closed, resulting in even larger sway variability and higher mean speeds.

Besides showing that the balance task manipulations were effective in creating marked differences in COP behavior, the repeated-measures ANOVA on the pre-intervention COP outcome measures revealed no significant differences between the experimental and control group in this regard. Hence, no relevant systematic differences in postural control were present between both groups before intervention.

### 4.4. Sleep quantity and quality

The mean rated sleep quantity of the participants in the experimental group was 7.5 ± 0.9 before both the familiarization and intervention session, while that of the participants in the control group was 7.3 ± 1.0 before their intervention session. The mean rating of the sleep quality of the participants in the experimental group was 1.8 ± 0.8 before the familiarization session and 2.0 ± 0.6 before the intervention session, while that of the participants in the control group was 1.9 ± 0.5 before their intervention session. Mann-Whitney tests revealed no significant differences between both groups regarding sleep quantity, *U* = 191.00, *p* = 0.32, *r* = -0.15, and quality, *U* = 261.00, *p* = 0.39, *r* = -0.13, before the interventions nor between the familiarization and intervention sessions of the experimental group: sleep quantity, *U* = 208.50, *p* = 0.76, *r* = -0.05, sleep quality, *U* = 254.50, *p* = 0.32, *r* = 0.16.

### 4.5. Subjective levels of induced MF

The subjective levels of MF induced by the interventions are shown in Fig. 3. Following the interventions significantly higher absolute levels of MF were reported by the experimental group (*M* = 70.54, *SE* = 2.51) than the control group (*M* = 41.59, *SE* = 4.77), *U* = 53.00, *p* < 0.001, *r* = -0.66. The relative subjective level of MF was rated significantly higher by the experimental group (*M* = 25.20, *SE* = 2.88) than the control group (*M* = 0.05, *SE* = 2.43), *U* = 33.00, *p* < 0.001, *r* = -0.73. These results indicate that the TloadDback task was effective in inducing MF and that watching the documentary did not affect mental fatigue levels.

**Fig. 3.**
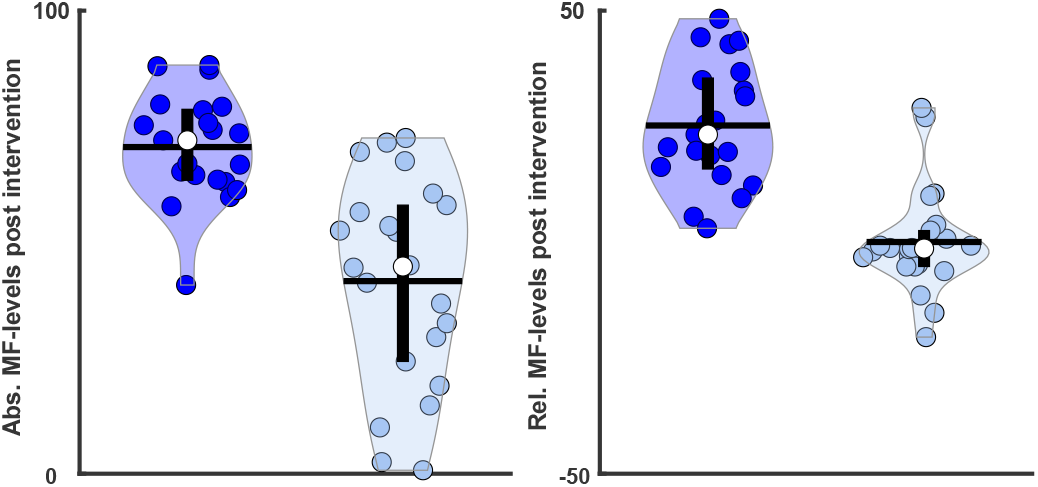
Violin plots of subjective levels of induced MF; Left panel: absolute subjective level of MF after the interventions (TloadDback [dark blue] vs. documentary [light blue]); Right panel: relative subjective level of MF after compared to before the intervention. Each dot represents one participant, and the size of the violin the spread of the data. Horizontal bar represents the mean; vertical bar is the interquartile range, and the white center dot is the median.

### 4.6. Influence of MF on COP behavior

Violin plots of the COP relative change scores are shown in Fig. 4 and 5, allowing a qualitative visual inspection of the intervention results. The results of the 2 x 2 x 3 repeated-measures ANOVAs on the relative change scores are collected in Tables 3 and 4, allowing a quantitative assessment. No significant group effects were present for any of the three COP outcome measures. Moreover, *t*-tests revealed that the relative change scores of the COP outcome measures did not differ significantly from a test value of zero, i.e., over all tasks in both directions (see Table A.1 and 2). The absence of significant main effects of posture and condition implies that the balance tasks were performed in a similar manner before and after the interventions.

**Table 3.** Statistical results of the repeated-measures ANOVA on the relative change of sway variability and mean speed values; *df* = degree of freedom, * = significant

| Effects | | $\Delta$ Sway variability | | | | $\Delta$ Mean speed | | | |
| --- | --- | --- | --- | --- | --- | --- | --- | --- | --- |
| | | *df* (1,2) | *F* | *p* | $\eta_p^2$ | *df* (1,2) | *F* | *p* | $\eta_p^2$ |
| Group | ML | 1,41 | 2.60 | 0.115 | 0.06 | 1,41 | 0.57 | 0.455 | 0.01 |
|  | AP | 1,41 | 0.23 | 0.637 | 0.01 | 1,41 | 2.01 | 0.164 | 0.05 |
| Posture | ML | 1,41 | 1.44 | 0.238 | 0.03 | 1,41 | 0.36 | 0.553 | 0.01 |
|  | AP | 1,41 | 0.89 | 0.352 | 0.02 | 1,41 | 2.54 | 0.118 | 0.06 |
| Condition | ML | 1.8,75.8 | 0.20 | 0.803 | 0.01 | 1.7,70.5 | 1.33 | 0.270 | 0.03 |
|  | AP | 2.0,80.2 | 0.01 | 0.992 | 0.00 | 1.9,76.2 | 1.52 | 0.225 | 0.04 |
| Posture x Group | ML | 1,41 | 5.23 | **0.027\*** | 0.11 | 1,41 | 0.07 | 0.796 | 0.00 |
|  | AP | 1,41 | 1.82 | 0.185 | 0.04 | 1,41 | 0.20 | 0.657 | 0.01 |
| Condition x Group | ML | 1.8,75.8 | 2.53 | 0.091 | 0.06 | 1.7,70.5 | 1.90 | 0.163 | 0.04 |
|  | AP | 2.0,80.2 | 0.01 | 0.203 | 0.04 | 1.9,76.2 | 3.02 | 0.058 | 0.07 |
| Posture x | ML | 1.9,78.9 | 1.64 | 0.202 | 0.04 | 1.9,79.2 | 0.99 | 0.357 | 0.02 |
| Condition | AP | 2.0,80.6 | 0.61 | 0.546 | 0.02 | 1.8,73.2 | 0.00 | 0.999 | 0.00 |
| Posture x | ML | 1.9,78.9 | 0.74 | 0.477 | 0.02 | 1.9,79.2 | 2.28 | 0.111 | 0.05 |
| Condition x Group | AP | 2.0,80.6 | 0.69 | 0.503 | 0.02 | 1.8,73.2 | 0.31 | 0.702 | 0.01 |

**Table 4.** Statistical results of the repeated-measures ANOVA on the relative change of sample entropy values; left: calculated using Lake et al.’s method; right: calculated using Montesinos et al.’s method; *df* = degree of freedom, * = significant

| | | $\Delta$ Sample entropy: $m = 3, r = 0.05$ | | | | $\Delta$ Sample entropy: $m = 4, r = 0.25$ | | | |
| --- | --- | --- | --- | --- | --- | --- | --- | --- | --- |
| Effects | | *df* (1,2) | *F* | *p* | $\eta_p^2$ | *df* (1,2) | *F* | *p* | $\eta_p^2$ |
| Group | ML | 1,41 | 1.09 | 0.304 | 0.03 | 1,41 | 1.38 | 0.247 | 0.03 |
|  | AP | 1,41 | 0.11 | 0.742 | 0.00 | 1,41 | 0.03 | 0.862 | 0.01 |
| Posture | ML | 1,41 | 1.04 | 0.314 | 0.03 | 1,41 | 0.18 | 0.676 | 0.00 |
|  | AP | 1,41 | 3.05 | 0.088 | 0.07 | 1,41 | 2.00 | 0.165 | 0.05 |
| Condition | ML | 1.8,75.2 | 0.72 | 0.477 | 0.02 | 1.9,76.2 | 0.39 | 0.671 | 0.01 |
|  | AP | 2.0,80.9 | 0.08 | 0.918 | 0.00 | 2.0,80.6 | 0.15 | 0.86 | 0.00 |
| Posture x Group | ML | 1,41 | 4.17 | **0.048\*** | 0.09 | 1,41 | 3.35 | 0.075 | 0.08 |
|  | AP | 1,41 | 1.24 | 0.272 | 0.03 | 1,41 | 1.41 | 0.241 | 0.03 |
| Condition x Group | ML | 1.8,75.2 | 3.28 | **0.047\*** | 0.07 | 1.9,76.2 | 2.66 | 0.078 | 0.06 |
|  | AP | 2.0,80.9 | 3.77 | **0.028\*** | 0.08 | 2.0,80.6 | 2.95 | 0.059 | 0.07 |
| Posture x | ML | 1.9,76.9 | 1.78 | 0.178 | 0.04 | 1.9,79.5 | 1.57 | 0.215 | 0.04 |
| Condition | AP | 1.9,79.5 | 0.83 | 0.437 | 0.02 | 2.9,77.0 | 1.24 | 0.293 | 0.03 |
| Posture x | ML | 1.9,76.9 | 0.79 | 0.451 | 0.02 | 1.9,79.5 | 1.29 | 0.280 | 0.03 |
| Condition x Group | AP | 1.9,79.5 | 1.84 | 0.167 | 0.04 | 2.9,77.0 | 0.53 | 0.578 | 0.01 |

**Fig. 4.**
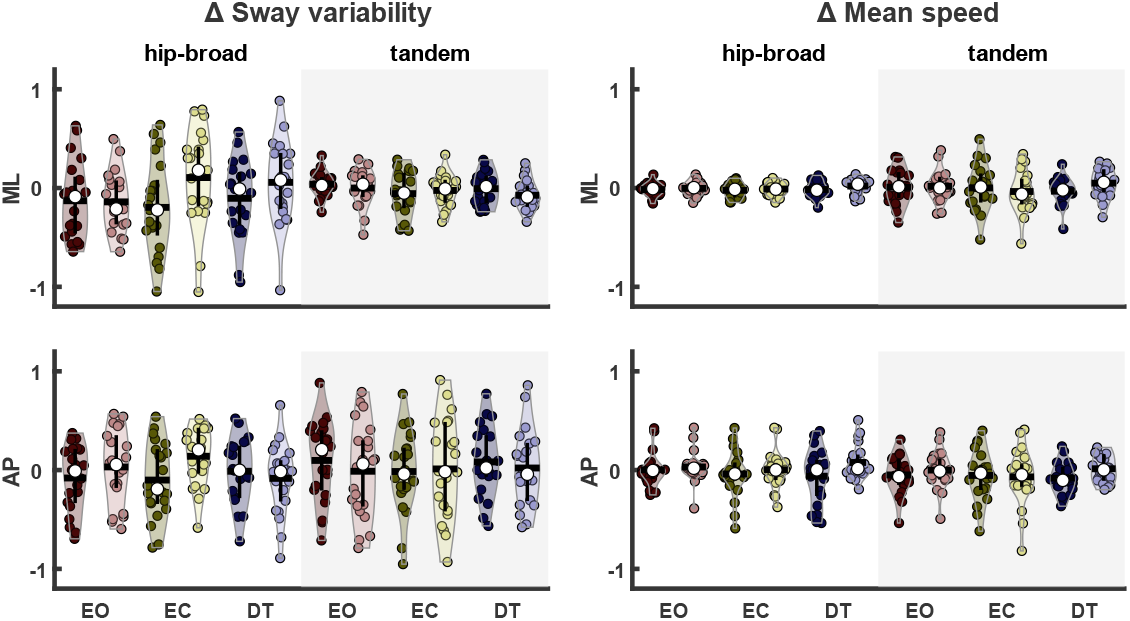
Left panels: relative change of sway variability; right panels: relative change of mean speed in ML (above) and AP (below) direction. Experimental group [dark colors], control group [light colors]. Each dot represents one participant, and the violin the spread of the data. Horizontal bar represents the mean; vertical bar is the interquartile range, and the white center dot is the median.

**Fig. 5.**
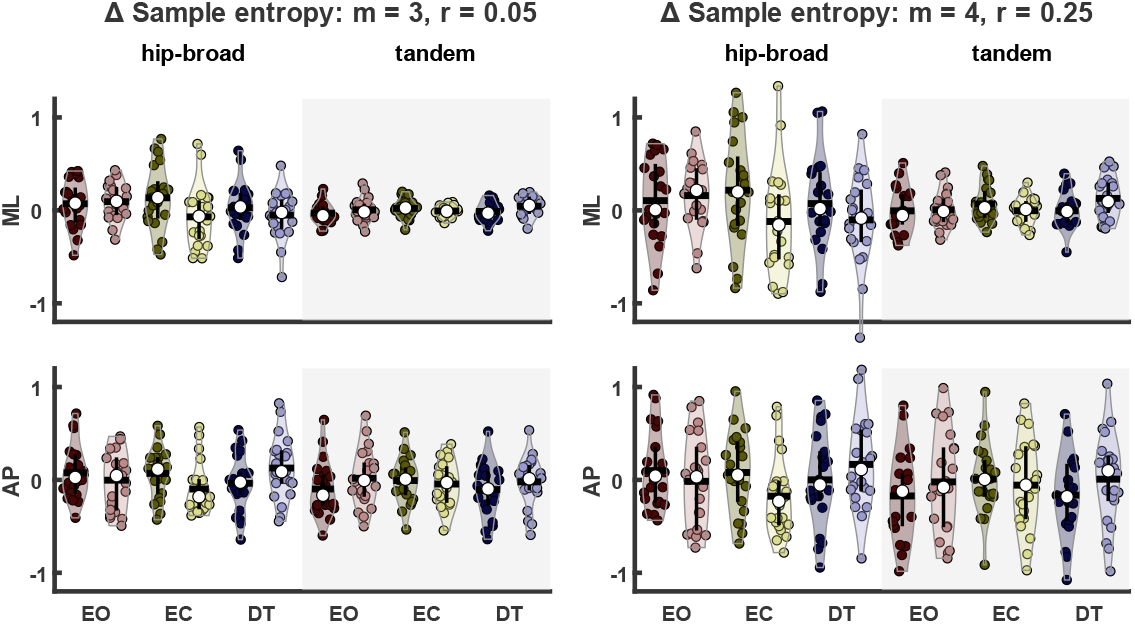
Left panels: relative change of sample entropy, according to Lake et al.’s method; right panels: relative change of sample entropy according to Montesinos et al.’s method in ML (above) and AP (below) direction. Experimental group [dark colors], control group [light colors]. Each dot represents one participant, and the violin the spread of the data. Horizontal bar represents the mean; vertical bar is the interquartile range, and the white center dot is the median.

Although no significant main effects occurred, a few weakly significant interaction effects were observed. Significant group x posture interaction effects were found for both sway variability and sample entropy using Lake et al.’s method in the ML-direction, but not in the AP-direction. These effects occurred because in the control group neither sway variability nor sample entropy changed due to the intervention in either posture, whereas in the experimental group a decrease of sway variability and an increase of sample entropy occurred in the hip-broad stance, but not in the tandem stance. Additionally, significant condition x group interactions were found for the relative change in sample entropy using Lake et al.’s settings in both sway directions. Post-hoc tests revealed that in the eyes-closed condition sample entropy values were affected differently by the two interventions. Whereas the relative change in sample entropy was positive for the experimental group, indicating less regularity and greater automaticity, the relative change was negative for the control group, indicating greater regularity and less automaticity. These significant interactions were absent when using Montesinos et al.’s method for calculating the sample entropy.

## 5. Discussion

This study was conducted to examine the effect of MF on postural control during quiet standing in young adults. We ventured to do so in a rigorous and encompassing manner to help resolve the existing ambiguity in pertinent studies on this research topic. In line with the prevailing assumption in previous studies on the issue of interest, we hypothesized that, in young adults, MF leads to a worsening of the ability to maintain quiet upright stance, resulting in increased COP sway variability and mean speed, and lower sample entropy values. Additionally, we hypothesized that the influence of mental fatigue on balance control depends on the attentional effort required by the balance tasks being performed. To test these hypotheses, we included several static balance tasks in the experimental design that differed in difficulty and attentional demands, which were performed both before and after the interventions to assess the effect of MF.

### 5.1. COP behavior before intervention

Before testing the main research hypotheses regarding the effect of MF on balance control, we verified that the six balance tasks varied in difficulty and attentional demands before the intervention by comparing the selected COP outcome measures across tasks. This was the case. The COP time-series differed significantly between the two postures as well as between the three conditions in which these postures were maintained (eyes open, eyes closed, and dual-task). The effect of posture on the three COP outcome measures was clear-cut and confirmed our expectation that standing in tandem stance is less stable than standing in hip-broad stance.

Furthermore, post-hoc analyses confirmed our expectation that standing with eyes closed is less stable than standing with eyes open (as evidenced by a significant increase in postural sway and mean speed), and that the introduction of a cognitive dual-task reduces attentional control over the balance task being performed (as evidenced by a significant increase in sample entropy). Although not all post-hoc comparisons between the balance conditions were significant for the three COP outcome measures, the significant effects were all in the expected direction. The effect of condition on the sample entropy measures was less pronounced than expected (e.g., sample entropy values were not significantly lower for standing with eyes closed compared to standing with eyes open). This was also evident from the fact that the posture x condition interaction was significant for sway variability and mean speed, but not for sample entropy, regardless of the method used for its calculation. Sample entropy is a sensitive measure that may easily be affected by design features or peculiarities of the time-series obtained.

Nevertheless, the obtained findings are in line with previous studies which have shown that: (i) a posture with a smaller base-of-support is more challenging (Lee & Shin, 2019; Sarabon, Rosker, Loefler, & Kern, 2013), (ii) postural control becomes more ‘automatic’ and efficient when attention is directed externally, i.e. with eyes open (Potvin-Desrochers, et al., 2017; Rhea, et al., 2019; Richer & Lajoie, 2020; J. F. Stins, et al., 2009; Yamada & Raisbeck, 2021) and less ‘automatic’ and efficient when attention is directed internally, i.e. with eyes closed (Becker & Hung, 2020; Donker, et al., 2007; J. F. Stins, et al., 2009), and that cognitive tasks promote automatization of postural control in young and older adults (Donker, et al., 2007).

Importantly, the analysis of the COP behavior before intervention not only showed that the balance task manipulation was effective, but also that no significant difference existed in this regard between the groups, implying that any significant group difference in COP behavior after the intervention could be attributed to the intervention.

### 5.2. Sleep quantity and quality

We verified that no significant differences existed in the quantity and quality of sleep that the experimental and control group had received before the intervention since any differences in this regard could confound the experimental results. For the same reason, we verified that no significant differences existed in the quantity and quality of sleep of the experimental group before the extra session and before the intervertion. Hence, any group effect could not attributed to a difference in sleep, neither directly nor indirectly (through a difference in fatiguability due to sleep of the experimental group between the familiarization session and the intervention).

### 5.3. Subjective levels of induced MF

As a final preliminary step before addressing the main research hypotheses, we verified that the method we used to induce fatigue in an individually tailored manner (i.e., adapted to an individually predefined maximum cognitive load) was effective, and led to significantly higher subjective levels of MF than watching a documentary. The experimental group reported significantly higher levels of subjective MF after performing the TloadDback task with individualized SDT, compared to the control group. Furthermore, the individualized TloadDback task led to comparable subjective levels of MF reported by the participants of the experimental group, as indicated by the small between-subject variance (Borragan, et al., 2017). In contrast, the variation of subjective levels of MF after the intervention reported by the participants of the control group was considerable (see Fig. 3, left).

However, when participants were asked to reflect about the change in their MF level, by comparing their MF status before and after the intervention, the participants of the control group stated not to have experienced any change in MF (see Fig. 3, right). One possible explanation of this paradoxical finding is that the MF levels of the control subjects were already broadly distributed before the intervention and left unaffected by watching the documentary. However, this is impossible to verify as we deliberately refrained from assessing subjective levels of MF before the intervention. Another explanation is that the broad variation in post-intervention MF ratings is a genuine reflection of how the control participants felt after having watched the documentary, but somehow could or did not relate this state to how they felt before, perhaps as a result of cognitive considerations. The verbal feedback provided by the control participants after the experiment speaks in favor of this second explanation: some described the documentary as ‘boring’, some said it made them feel ‘relaxed’ or ‘sleepy’, while others qualified the documentary as interesting and stimulating. This discussion illustrates that finding an adequate control intervention in experimental studies on the influence of MF on task performance remains difficult. The documentary used in the present study was chosen because it was also used in comparable studies by Hachard, et al. (2020) and (Varas-Diaz, et al., 2020), but like any other documentary, may well have the drawback some participants find it entertaining and other participants less so or not at all.

### 5.4. Influence of MF on postural control

By verifying the effectiveness of the balance task manipulation and the chosen method to induce similar subjective levels of MF, and by ruling out sleep quantity and quality as possible confounds, we fulfilled the necessary conditions for testing the research hypotheses. Overall, the research hypotheses were disconfirmed: no significant main effects of group were found on any of the COP measures of interest, indicating that the experimentally induced MF, although clearly present in the experimental group, had no statistically discernable effect on static balance control in the population of interest. In the absence of a main effect of group, also the hypothesis that the influence of MF on balance control depends on the attentional effort required by the balance tasks being performed was generally disconfirmed. The only statistical evidence found in favor of this hypothesis were a few weakly significant interaction effects involving the factor group. A significant posture x group interaction effect was found in the ML-direction for the relative change in sway variability and sample entropy when calculated according to Lake et al.’s method, but not when calculated according to Montesinos et al.’s method. This effect occurred because MF affected postural control in the ML direction in hip-broad stance but not in tandem stance. This finding precludes a clear interpretation, since an effect on tandem stance rather than hip-broad stance would have been expected, assuming MF hampers balance control through depletion of cognitive resources. Additionally, significant condition x group interactions were found for the relative change in sample entropy when calculated according to Lake et al.’s method, but not when calculated according to Montesinos et al.’s method. These effects occurred because the relative change score of sample entropy was positive in the eyes-closed condition in the experimental group and negative in the control group and could thus at least partly be interpreted in terms of a depletion of cognitive resources (resulting in less attentional control and thus less regularity and greater automaticity). However, this finding was non-robust as it proved to be critically dependent on the precise settings for calculating sample entropy. Collectively, these results suggest that, in young adults, postural control in static balance tasks is largely automatic and only marginally affected by MF.

One could argue that the level of MF induced by the TloadDback task, was not severe enough, even though participants in the experimental group indicated clearly that they felt more fatigued after having performed this task. Whereas Borragan, et al. (2017) and O’Keeffe, et al. (2019) showed that performing the TloadDback task for 16 minutes is sufficient to mentally fatigue participants to a sufficiently high degree, Jacquet, Poulin-Charronnat, Bard, and Lepers (2020) argued that the standardized duration of the TloadDback task of 16 minutes is too short to affect physical performance. According to them, performing a mentally fatiguing task for such a short duration intervenes primarily with people’s perception of fatigue, rather than inducing sufficient MF to reduce their motor abilities. Pitts and Bhatt (2023) raised a similar concern. Although this concern is certainly valid, it should not be used to dismiss any null finding as invalid, certainly not in situations, such as in the present study, where the attentional component of balance control is expected to be small. Deschamps, et al. (2013) and Hachard, et al. (2020) used mentally fatiguing tasks of 30- and 90-minute duration, respectively, but also found no (or no strong) effects of MF. Deschamps, et al. (2013) found no effects of MF while standing on a stable surface with eyes open or closed; only while standing on an unstable surface effects of MF became manifest. Hachard, et al. (2020) not only found a reduced efficiency of postural control in different conditions after having mentally fatigued half of the participants, but also in the other half of the participants, which served as control group. It therefore seemed that in this case the observed changes in postural control did not result from the experimentally induced MF, but from some confounding variable (e.g., sitting). A significant effect on postural control between the experimental and control group was only observed during standing with eyes closed on a stable surface. The results of Deschamps, et al. (2013) and Hachard, et al. (2020) thus indicate that the effect of MF on postural control does not become stronger nor more obvious for longer durations of the mentally fatiguing task.

Since no consistent effects of MF on postural control were found across studies, it might be that, in healthy young adults, static balance does not require sufficient attentional resources to be significantly affected by the cognitive resource depletion resulting from experimentally induced MF. Despite the evidence that even simple static balance tasks require investment of attentional resources, maintaining postural stability during unperturbed upright standing is widely regarded as a largely automatic and reflex-based process, which demands only very little attention (Kerr, et al., 1985; Teasdale, et al., 1993). This line of thinking is supported by the observation that older adults seem more sensitive to MF. In a study by Varas-Diaz, et al. (2020), with a design similar to our study, significant impacts of MF in healthy older adults as well as people who had suffered a stroke were observed. Postural control was found to be impaired under various sensory conditions and when concurrently performing a cognitive task. These enhanced effects of MF might derive from the fact that attentional demands to maintain equilibrium during static balance tasks increase with aging and pathological conditions (Bisson, McEwen, Lajoie, & Bilodeau, 2011; Teasdale, et al., 1993). In contrast to healthy young adults for whom the performance of simple static balance tasks seems effortless, neurodegenerative processes reduce postural control in older adults, rendering balance maintenance more difficult (Bergamin, et al., 2014). This is reflected in increased sway and decreased sample entropy values (Roerdink, et al., 2006). Apart from physical changes, the increased effort for postural control can also be explained by reduced cognitive functions (Amboni, Barone, & Hausdorff, 2013). We conclude that the chosen postural tasks in this experiment required too little attention and effort such that the evoked depletion of cognitive resources did not interfere with postural control in our sample of healthy young adults.

Future studies assessing whether (and if so, which) postural parameters are affected by experimentally induced MF should either make the postural tasks more challenging and thus attention demanding (Lew & Qu, 2014), or intensify the level of induced MF, which might be achieved by sleep deprivation for a prolonged time (Batuk, Batuk, & Aksoy, 2020; Cheng, et al., 2018; Patel, et al., 2008).

## 6. Strengths and limitations

The study has two main strengths. First, due to the comparatively large sample size of participants the study is well-powered, rendering its results robust and meaningful. Second, the inter-individual variation in the level of induced MF was minimized by individualizing the stimulus duration time of the TloadDback task. The study also has two main limitations. First, because the level of MF induced by the intervention was rated only subjectively, it cannot be ruled out that the subjective ratings were influenced by other confounding factors, such as perceptual, memory, and affective processes. Second, and relatedly, the absolute and relative subjective ratings of MF by the control group (after having watched the documentary) were paradoxical and not readily interpretable, rendering it not fully certain that watching the documentary induced no MF whatsoever.

## 7. Conclusion

Based on the premise that cognitive, attentional control plays a significant role in the performance of balance tasks, it is often assumed that mental fatigue impedes balance by depleting cognitive resources. However, in healthy young adults the influence of mental fatigue and the associated depletion of cognitive resources on the performance of static balance tasks is marginal at best. In this population and task domain, postural control is largely automatic with minimal cognitive mediation.

## Data and code availability

To allow reproducibility of our results and any further analyses all data and code can be found here: https://osf.io/jatc6/.

## Declarations

### Ethical approval

This study was approved by the local ethics committee of the Faculty of Behavioral and Movement Sciences of the Vrije Universiteit Amsterdam (VCWE-2022-154R2)

## Appendix A. Appendix

**Table A. 1.** T-test of sway variability and mean speed against a test-value of zero; H = hip broad stance, T = tandem stance, EO = eyes open, EC = eyes closed, DT = dual task

| Tasks | | $\Delta$ Sway variability | | | | $\Delta$ Mean speed | | | |
| --- | --- | --- | --- | --- | --- | --- | --- | --- | --- |
|  |  | Experiment |  | Control |  | Experiment |  | Control |  |
|  |  | *t* | *sig* | *t* | *sig* | *t* | *sig* | *t* | *sig* |
| H-EO | ML | -1.53 | 0.14 | -2.13 | 0.05 | -1.08 | 0.29 | 0.02 | 0.98 |
|  | AP | -1.16 | 0.26 | 0.40 | 0.69 | 0.08 | 0.94 | 0.94 | 0.36 |
| H-EC | ML | -1.93 | 0.07 | 1.03 | 0.31 | -1.44 | 0.17 | 0.05 | 0.96 |
|  | AP | -1.25 | 0.23 | 2.25 | 0.04 | -0.80 | 0.43 | 0.08 | 0.94 |
| H-DT | ML | -1.21 | 0.24 | 0.76 | 0.45 | -0.87 | 0.40 | -0.17 | 0.87 |
|  | AP | -0.17 | 0.86 | -1.13 | 0.27 | -1.06 | 0.30 | 2.11 | 0.06 |
| T-EO | ML | 1.43 | 0.17 | -0.11 | 0.92 | 0.22 | 0.83 | 1.14 | 0.27 |
|  | AP | 1.14 | 0.27 | -0.12 | 0.91 | -1.23 | 0.23 | 0.01 | 1.00 |
| T-EC | ML | -1.16 | 0.26 | -0.32 | 0.75 | 0.02 | 0.98 | -0.22 | 0.83 |
|  | AP | -0.17 | 0.87 | 0.11 | 0.92 | -1.03 | 0.32 | -1.16 | 0.26 |
| T-DT | ML | -0.19 | 0.85 | -2.40 | 0.04 | -1.09 | 0.29 | 1.23 | 0.23 |

**Table A. 2.** T-test of sample entropy change-score values against a test-value of zero; r right: parameters were chosen according to Montesinos et al., 2018; left: parameters were chosen according to Lake et al., 2002; H = hip broad stance, T = tandem stance, EO = eyes open, EC = eyes closed, DT = dual task

| Tasks | | $\Delta$ Sample entropy: m = 3, r = 0.05 | | | | $\Delta$ Sample entropy: m = 4, r = 0.25 | | | |
| --- | --- | --- | --- | --- | --- | --- | --- | --- | --- |
|  |  | Experiment |  | Control |  | Experiment |  | Control |  |
|  |  | *t* | *sig* | *t* | *sig* | *t* | *sig* | *t* | *sig* |
| H-EO | ML | 1.31 | 0.21 | 2.26 | 0.05 | 1.06 | 0.30 | 2.05 | 0.05 |
|  | AP | 1.26 | 0.22 | -0.05 | 0.96 | 1.15 | 0.26 | -0.15 | 0.89 |
| H-EC | ML | 1.79 | 0.09 | -0.97 | 0.35 | 1.71 | 0.10 | -0.94 | 0.36 |
|  | AP | 1.26 | 0.22 | -1.68 | 0.11 | 0.89 | 0.39 | -1.95 | 0.06 |
| H-DT | ML | 0.48 | 0.63 | -0.97 | 0.35 | 0.68 | 0.50 | -1.04 | 0.31 |
|  | AP | -0.57 | 0.57 | 1.81 | 0.09 | 0.01 | 1.00 | 1.62 | 0.12 |
| T-EO | ML | -1.25 | 0.23 | 0.59 | 0.56 | -0.10 | 0.92 | 0.49 | 0.63 |
|  | AP | -1.60 | 0.13 | 0.28 | 0.79 | -1.74 | 0.10 | -0.19 | 0.85 |
| T-EC | ML | 1.00 | 0.33 | -0.19 | 0.85 | 1.64 | 0.12 | 0.13 | 0.90 |
|  | AP | -0.14 | 0.89 | -0.88 | 0.39 | 0.11 | 0.92 | -0.56 | 0.58 |
| T-DT | ML | -0.53 | 0.60 | 2.06 | 0.06 | -0.11 | 0.91 | 2.60 | 0.03 |

## References

Amboni, M., Barone, P., & Hausdorff, J. M. (2013). Cognitive contributions to gait and falls: evidence and implications. Movement Disorders, 28, 1520–1533 https://doi.org/10.1002/mds.25674.

Barrouillet, P., Bernardin, S., Portrat, S., Vergauwe, E., & Camos, V. (2007). Time and cognitive load in working memory. J Exp Psychol Learn Mem Cogn, 33, 570–585 https://doi.org/10.1037/0278-7393.33.3.570.

Batuk, I. T., Batuk, M. O., & Aksoy, S. (2020). Evaluation of the postural balance and visual perception in young adults with acute sleep deprivation. J Vestib Res, 30, 383–391 https://doi.org/10.3233/VES-200778.

Becker, K. A., & Hung, C. J. (2020). Attentional focus influences sample entropy in a balancing task. Hum Mov Sci, 72, 102631 https://doi.org/10.1016/j.humov.2020.102631.

Behrens, M., Mau-Moeller, A., Lischke, A., Katlun, F., Gube, M., Zschorlich, V., Skripitz, R., & Weippert, M. (2017). Mental fatigue increases gait variability during dual-task walking in old adults. The Journals of Gerontology: Series A, 73, 792–797-792–797 https://doi.org/10.1093/gerona/glx210.

Bergamin, M., Gobbo, S., Zanotto, T., Sieverdes, J. C., Alberton, C. L., Zaccaria, M., & Ermolao, A. (2014). Influence of age on postural sway during different dual-task conditions. Front Aging Neurosci, 6, 271 https://doi.org/10.3389/fnagi.2014.00271.

Bisson, E. J., McEwen, D., Lajoie, Y., & Bilodeau, M. (2011). Effects of ankle and hip muscle fatigue on postural sway and attentional demands during unipedal stance. Gait Posture, 33, 83–87 https://doi.org/10.1016/j.gaitpost.2010.10.001.

Boksem, M. A., Meijman, T. F., & Lorist, M. M. (2005). Effects of mental fatigue on attention: an ERP study. Cognitive Brain Research, 25, 107–116 https://doi.org/10.1016/j.cogbrainres.2005.04.011.

Borragan, G., Slama, H., Bartolomei, M., & Peigneux, P. (2017). Cognitive fatigue: A Time-based Resource-sharing account. Cortex, 89, 71–84 https://doi.org/10.1016/j.cortex.2017.01.023.

Borragan, G., Slama, H., Destrebecqz, A., & Peigneux, P. (2016). Cognitive Fatigue Facilitates Procedural Sequence Learning. Front Hum Neurosci, 10, 86 https://doi.org/10.3389/fnhum.2016.00086.

Brahms, M., Heinzel, S., Rapp, M., Muckstein, M., Hortobagyi, T., Stelzel, C., & Granacher, U. (2022). The acute effects of mental fatigue on balance performance in healthy young and older adults - A systematic review and meta-analysis. Acta Psychol (Amst), 225, 103540 https://doi.org/10.1016/j.actpsy.2022.103540.

Burr, J., & Nesselroade, J. (1990). Statistical Methods in Longitudinal Research: Principles and Structuring Change (Vol. 1): American Press Inc.

Buysse, D. J., Reynolds, C. F., 3rd, Monk, T. H., Berman, S. R., & Kupfer, D. J. (1989). The Pittsburgh Sleep Quality Index: a new instrument for psychiatric practice and research. Psychiatry Res, 28, 193–213 https://doi.org/10.1016/0165-1781(89)90047-4.

Cheng, S., Ma, J., Sun, J., Wang, J., Xiao, X., Wang, Y., & Hu, W. (2018). Differences in sensory reweighting due to loss of visual and proprioceptive cues in postural stability support among sleep-deprived cadet pilots. Gait & Posture, 63, 97–103-197–103 https://doi.org/10.1016/j.gaitpost.2018.04.037.

Cohen, J. (1988). Statistical power analysis for the behavioral sciences: Lawrence Erlbaum Associates.

Deschamps, T., Beauchet, O., Annweiler, C., Cornu, C., & Mignardot, J. B. (2014). Postural control and cognitive decline in older adults: position versus velocity implicit motor strategy. Gait Posture, 39, 628–630 https://doi.org/10.1016/j.gaitpost.2013.07.001.

Deschamps, T., Magnard, J., & Cornu, C. (2013). Postural control as a function of time-of-day: influence of a prior strenuous running exercise or demanding sustained-attention task. J Neuroeng Rehabil, 10, 26 https://doi.org/10.1186/1743-0003-10-26.

Donker, S. F., Roerdink, M., Greven, A. J., & Beek, P. J. (2007). Regularity of center-of-pressure trajectories depends on the amount of attention invested in postural control. Exp Brain Res, 181, 1–11 https://doi.org/10.1007/s00221-007-0905-4.

Faul, F., Erdfelder, E., Lang, A. G., & Buchner, A. (2007). G*Power 3: a flexible statistical power analysis program for the social, behavioral, and biomedical sciences. Behav Res Methods, 39, 175–191 https://doi.org/10.3758/bf03193146.

Gorrall, B. K., Curtis, J., Little, T. D., & Panko, P. (2016). Innovations in Measurement: Visual Analog Scales and Retrospective Pretest Self-Report Designs. Actualidades en Psicología, 30, 1–6 https://doi.org/10.15517/ap.v30i120.22932.

Grobe, S., Kakar, R. S., Smith, M. L., Mehta, R., Baghurst, T., & Boolani, A. (2017). Impact of cognitive fatigue on gait and sway among older adults: A literature review. Preventive Medicine Reports, 6, 88–93-88–93 https://doi.org/10.1016/j.pmedr.2017.02.016.

Hachard, B., Noé, F., Ceyte, H., Trajin, B., & Paillard, T. (2020). Balance control is impaired by mental fatigue due to the fulfilment of a continuous cognitive task or by the watching of a documentary. Experimental Brain Research, 238, 861–868 https://doi.org/10.1007/s00221-020-05758-2.

Hockey, R. (2013). The psychology of fatigue; work effort and control: Cambridge University Press.

Jacquet, T., Poulin-Charronnat, B., Bard, P., & Lepers, R. (2020). Persistence of Mental Fatigue on Motor Control. Front Psychol, 11, 588253 https://doi.org/10.3389/fpsyg.2020.588253.

Kerr, B., Condon, S. M., & McDonald, L. A. (1985). Cognitive spatial processing and the regulation of posture. J Exp Psychol Hum Percept Perform, 11, 617–622 https://doi.org/10.1037/0096-1523.11.5.617.

Lajoie, Y., Teasdale, N., Bard, C., & Fleury, M. (1993). Attentional demands for static and dynamic equilibrium. Exp Brain Res, 97, 139–144 https://doi.org/10.1007/BF00228824.

Lake, D. E., Richman, J. S., Griffin, M. P., & Moorman, J. R. (2002). Sample entropy analysis of neonatal heart rate variability. Am J Physiol Regul Integr Comp Physiol, 283, R789–797 https://doi.org/10.1152/ajpregu.00069.2002.

Lee, Y., & Shin, S. (2019). Effects of the shape of the base of support and dual task execution on postural control. The Asian Journal of Kinesiology, 21, 14–24-14–24 https://doi.org/10.15758/ajk.2019.21.1.14.

Lew, F. L., & Qu, X. (2014). Effects of mental fatigue on biomechanics of slips. Ergonomics, 57, 1927–1932 https://doi.org/10.1080/00140139.2014.937771.

Martínez-Cagigal, V. (2018). Sample Entropy. In. Mathworks Montesinos, L., Castaldo, R., & Pecchia, L. (2018). On the use of approximate entropy and sample entropy with centre of pressure time-series. J Neuroeng Rehabil, 15, 116 https://doi.org/10.1186/s12984-018-0465-9.

Muir, S. W., Speechley, M., Wells, J., Borrie, M., Gopaul, K., & Montero-Odasso, M. (2012). Gait assessment in mild cognitive impairment and Alzheimer\textquotesingles disease: The effect of dual-task challenges across the cognitive spectrum. Gait & Posture, 35, 96–100-196–100 https://doi.org/10.1016/j.gaitpost.2011.08.014.

Nashner, L. M. (1976). Adapting reflexes controlling the human posture. Exp Brain Res, 26, 59–72 https://doi.org/10.1007/BF00235249.

Noé, F., Hachard, B., Ceyte, H., Bru, N., & Paillard, T. (2021). Relationship between the level of mental fatigue induced by a prolonged cognitive task and the degree of balance disturbance. Experimental Brain Research, 239, 2273–2283-2273–2283 https://doi.org/10.1007/s00221-021-06139-z.

O’Keeffe, K., Hodder, S., & Lloyd, A. (2019). A comparison of methods used for inducing mental fatigue in performance research: individualised, dual-task and short duration cognitive tests are most effective. Ergonomics, 63, 1–12-11–12 https://doi.org/10.1080/00140139.2019.1687940.

Papegaaij, S., Taube, W., Baudry, S., Otten, E., & Hortobagyi, T. (2014). Aging causes a reorganization of cortical and spinal control of posture. Front Aging Neurosci, 6, 28 https://doi.org/10.3389/fnagi.2014.00028.

Patel, M., Gomez, S., Berg, S., Almbladh, P., Lindblad, J., Petersen, H., Magnusson, M., Johansson, R., & Fransson, P. A. (2008). Effects of 24-h and 36-h sleep deprivation on human postural control and adaptation. Exp Brain Res, 185, 165–173 https://doi.org/10.1007/s00221-007-1143-5.

Pitts, J., & Bhatt, T. (2023). Effects of mentally induced fatigue on balance control: a systematic review. Exp Brain Res, 241, 13–30 https://doi.org/10.1007/s00221-022-06464-x.

Potvin-Desrochers, A., Richer, N., & Lajoie, Y. (2017). Cognitive tasks promote automatization of postural control in young and older adults. Gait Posture, 57, 40–45 https://doi.org/10.1016/j.gaitpost.2017.05.019.

Prado, J. M., Stoffregen, T. A., & Duarte, M. (2007). Postural sway during dual tasks in young and elderly adults. Gerontology, 53, 274–281 https://doi.org/10.1159/000102938.

Rhea, C. K., Diekfuss, J. A., Fairbrother, J. T., & Raisbeck, L. D. (2019). Postural control entropy is increased when adopting an external focus of attention. Motor Control, 23, 230–242-230–242 https://doi.org/10.1123/mc.2017-0089.

Richer, N., & Lajoie, Y. (2020). Automaticity of postural control while dual-tasking revealed in young and older adults. Experimental Aging Research, 46, 1–21-21–21 https://doi.org/10.1080/0361073X.2019.1693044.

Roerdink, M., De Haart, M., Daffertshofer, A., Donker, S. F., Geurts, A. C., & Beek, P. J. (2006). Dynamical structure of center-of-pressure trajectories in patients recovering from stroke. Exp Brain Res, 174, 256–269 https://doi.org/10.1007/s00221-006-0441-7.

Ruffieux, J., Keller, M., Lauber, B., & Taube, W. (2015). Changes in Standing and Walking Performance Under Dual-Task Conditions Across the Lifespan. Sports Med, 45, 1739–1758 https://doi.org/10.1007/s40279-015-0369-9.

Sarabon, N., Rosker, J., Loefler, S., & Kern, H. (2013). The effect of vision elimination during quiet stance tasks with different feet positions. Gait Posture, 38, 708–711 https://doi.org/10.1016/j.gaitpost.2013.03.005.

Schouppe, S., Danneels, L., Damme, S. V., Oosterwijck, S. V., Palmans, T., & Oosterwijck, J. V. (2019). Physical and cognitive exertion do not influence feedforward activation of the trunk muscles: a randomized crossover trial. Experimental Brain Research, 237, 3011–3021-3011–3021 https://doi.org/10.1007/s00221-019-05585-0.

Stins, J. F., Michielsen, M. E., Roerdink, M., & Beek, P. J. (2009). Sway regularity reflects attentional involvement in postural control: effects of expertise, vision and cognition. Gait Posture, 30, 106–109 https://doi.org/10.1016/j.gaitpost.2009.04.001.

Stins, J. F., Roerdink, M., & Beek, P. J. (2011). To freeze or not to freeze? Affective and cognitive perturbations have markedly different effects on postural control. Human Movement Science, 30, 190–202 https://doi.org/10.1016/j.humov.2010.05.013.

Takakusaki, K., Saitoh, K., Harada, H., & Kashiwayanagi, M. (2004). Role of basal ganglia-brainstem pathways in the control of motor behaviors. Neurosci Res, 50, 137–151 https://doi.org/10.1016/j.neures.2004.06.015.

Tanaka, M. (2015). Effects of mental fatigue on brain activity and cognitive performance: a magnetoencephalography study. Anatomy & Physiology, s4 https://doi.org/10.4172/2161-0940.s4-002.

Tassignon, B., Verschueren, J., Pauw, K. D., Roelands, B., Cutsem, J. V., Verhagen, E., & Meeusen, R. (2020). Mental fatigue impairs clinician-friendly balance test performance and brain activity. Translational Sports Medicine, 3, 616–625-616–625 https://doi.org/10.1002/tsm2.177.

Teasdale, N., Bard, C., LaRue, J., & Fleury, M. (1993). On the cognitive penetrability of posture control. Exp Aging Res, 19, 1–13 https://doi.org/10.1080/03610739308253919.

Teo, W. P., Goodwill, A. M., Hendy, A. M., Muthalib, M., & Macpherson, H. (2018). Sensory manipulation results in increased dorsolateral prefrontal cortex activation during static postural balance in sedentary older adults: An fNIRS study. Brain Behav, 8, e01109 https://doi.org/10.1002/brb3.1109.

Van Cutsem, J., Marcora, S., De Pauw, K., Bailey, S., Meeusen, R., & Roelands, B. (2017). The Effects of Mental Fatigue on Physical Performance: A Systematic Review. Sports Med, 47, 1569–1588 https://doi.org/10.1007/s40279-016-0672-0.

Varas-Diaz, G., Kannan, L., & Bhatt, T. (2020). Effect of Mental Fatigue on Postural Sway in Healthy Older Adults and Stroke Populations. Brain Sci, 10 https://doi.org/10.3390/brainsci10060388.

Vartia, Y. O. (1976). Relative changes and index numbers: The Research Institute of the Finnish Economy.

Verschueren, J. O., Tassignon, B., Proost, M., Teugels, A., Van Cutsem, J., Roelands, B., Verhagen, E., & Meeusen, R. (2020). Does mental fatigue negatively affect outcomes of functional performance tests? Medicine and science in sports and exercise, 52, 2002–2010 https://doi.org/10.1249/MSS.0000000000002323.

Yamada, M., & Raisbeck, L. D. (2021). The autonomy and focus of attention strategies under distraction: Frequency and sample entropy analyses in a dynamic balance task. Hum Mov Sci, 80, 102882 https://doi.org/10.1016/j.humov.2021.102882.

